# HIGH RESOLUTION ANNOTATION OF ZEBRAFISH TRANSCRIPTOME USING LONG-READ SEQUENCING

**DOI:** 10.1101/174821

**Authors:** German Nudelman, Antonio Frasca, Brandon Kent, Kirsten Edepli-Sadler, Stuart C. Sealfon, Martin J. Walsh, Elena Zaslavsky

**Affiliations:** Department of Neurology, Icahn School of Medicine at Mount Sinai, New York, NY, USA; Center for Advanced Research on Diagnostic Assays (CARDA); Department of Pharmacological Sciences, Icahn School of Medicine at Mount Sinai, New York, NY, USA; Department of Development and Regenerative Biology, Icahn School of Medicine at Mount Sinai, New York, NY, USA; Program in Biology, New York University Abu Dhabi, Abu Dhabi, UAE; Department of Genetics and Genomic Sciences, Icahn School of Medicine at Mount Sinai, New York, NY, USA

**Author notes:** These authors are joint first authors and contributed equally to this work (German Nudelman, Antonio Frasca).

## Abstract

With the emergence of zebrafish as an important model organism, a concerted effort has been made to study its transcriptome. This effort is limited, however, by gaps in zebrafish annotation, which are especially pronounced concerning transcripts dynamically expressed during zygotic genome activation (ZGA). To date, short read sequencing has been the principal technology for zebrafish transcriptome annotation. In part because these sequence reads are too short for assembly methods to resolve the full complexity of the transcriptome, the current annotation is rudimentary. By providing direct observation of full-length transcripts, recently refined long-read sequencing platforms can dramatically improve annotation coverage and accuracy. Here, we leveraged the SMRT platform to study transcriptome of zebrafish embryos before and after ZGA. Our analysis revealed additional novelty and complexity in the zebrafish transcriptome, identifying 2748 high confidence novel transcripts that originated from previously unannotated loci and 1835 high confidence new isoforms in previously annotated genes. We validated these findings using a suite of computational approaches including structural prediction, sequence homology and functional conservation analyses, as well as by confirmatory transcript quantification with short-read sequencing data. Our analyses provided insight into new homologs and paralogs of functionally important proteins and non-coding RNAs, isoform switching occurrences and different classes of novel splicing events. Several novel isoforms representing distinct splicing events were validated through PCR experiments, including the discovery and validation of a novel 8 kb transcript spanning multiple miR-430 elements, an important driver of early development. Our study provides a significantly improved zebrafish transcriptome annotation resource.

## INTRODUCTION

Characterization of the transcriptome provides insight into functional, physiological and biosynthetic cellular states. High-throughput short read sequencing (RNA-seq) has revolutionized study of the transcriptome (Ozsolak and Milos 2011), but is limited by the accuracy and completeness of the reference sequence annotation. Zebrafish is an important model organism and has been extensively used to study embryonic development (Howe et al. 2013). The first stage of zebrafish development is characterized by rapid cleavage of the embryonic cells and is entirely driven by the maternally provided mRNA and proteins as the zygotic genome is transcriptionally inactive until about 3 hours post-fertilization (3 hpf) when a cadre of developmentally essential genes become transcribed during zygotic genome activation (ZGA). Many studies have defined the transcriptome of the early embryo during these key developmental events and have identified thousands of genes that are induced during ZGA (Aanes et al. 2011; Pauli et al. 2012; Aanes et al. 2014; Heyn et al. 2014). The substantial maternal contribution provides for a relatively late onset of ZGA compared to other vertebrates, and makes zebrafish an ideal system to study this process.

Despite the significant advances in understanding the patterns of gene expression during ZGA, progress has been hampered by gaps in the annotation of the zebrafish genome and transcriptome. Recent studies profiling the zebrafish transcriptome using short-read RNA-sequencing technology during early embryogenesis suggest that thousands of transcripts are missing from the reference annotation (Aanes et al. 2011; Pauli et al. 2012; Aanes et al. 2014; Heyn et al. 2014, Wang et al. 2016b). In part, this can be attributed to the use of short read sequencing as the principal technology for understanding developmentally-regulated genes expressed in the early embryo and for zebrafish transcriptome annotation (Pauli et al. 2012; Pruitt et al. 2012; Heyn et al. 2014; Wang et al. 2016b). Due to its inherent length limitations, short read sequencing often fails to resolve transcriptional complexity associated with the mechanisms of alternative splicing, alternative transcription initiation, or alternative transcription termination sites that are ubiquitously utilized by eukaryotic organisms (Hiller et al. 2009; Florea and Salzberg 2013; Hooper 2014; Wang et al. 2016a). Further hampering the ability to completely annotate the zebrafish transcriptome using short-read data is a whole-genome duplication event, called the teleost-specific genome duplication (Postlethwait et al. 1998), which likely gave rise to sets of highly similar transcripts that are difficult to resolve without observing their full-length sequences.

Long-read sequencing can help overcome these limitations by providing direct observation of full-length transcripts. While earlier generations of long-read sequencing technologies were plagued by high sequencing error rates, technological and computational improvements have reduced the effective error rate to less than 1% (Rhoads and Au 2015). Due to its improved ability to define novel transcribed regions and disambiguate between distinct, but highly similar isoforms of annotated genes, long read-sequencing has become an important tool in the efforts to improve the transcriptome and genome annotation of several organisms (Sharon et al. 2013; Kim et al. 2014; Thomas et al. 2014; Gordon et al. 2015; Wang et al. 2016b, Abdel-Ghany et al. 2016).

In this study, we have utilized the current SMRT platform of Pacific Biosciences, which provides high sequencing accuracy and has no inherent upper limit on transcript length (Roberts et al. 2013; Rhoads and Au 2015), to improve annotation of the early ZGA-stage zebrafish transcriptome (Baroux et al. 2008; Tadros and Lipshitz 2009). Comparing our results with the most recent reference annotation (GRCz10) (Howe et al. 2013) revealed additional novelty and complexity in the early zebrafish transcriptome. We have identified thousands of novel transcripts from either annotated genes and previously unannotated loci. We validated many of these observations by comparison with short-read sequencing data as well as structural, homology and functional conservation analyses. We confirmed novel isoforms spanning key genes in early development, and extended observations of isoform switching pre- and post-ZGA. In particular, we identified a novel 8kb transcript spanning most miR-430 elements, which we validated using RT-qPCR. These transcripts were absent in the pre-ZGA samples, suggesting that they are among the earliest zygotically-expressed RNAs. In aggregate, our study provides an important new resource for zebrafish biologists with a much expanded and validated annotation of the zebrafish transcriptome.

## RESULTS

### Experimental system and long-read data characteristics

To characterize ZGA and maternal transcripts, we followed an experimental and computational pipeline illustrated in Fig. 1A. Embryos, generated from either the AB or hybrid Tubigen-AB line by natural spawning, were staged based on morphological criteria (Kimmel et al. 1995). Samples were collected in the presence and absence of transcriptional inhibition with α-amanitin, which blocks global transcription and specifically abrogates zygotic transcripts (Bensaude 2011; Tani and Akimitsu 2012). This approach was followed to help resolve a revised transcriptome with novel genes and isoforms derived from the early zebrafish zygote. Total RNA was extracted from pools of 15 embryos in each condition, as described (Bensaude 2011) and the cDNA libraries were sequenced on the PacBio SMRT platform. We also generated a short-read RNA-seq dataset collected from RNA of embryos at early and late stages in embryonic development.

**Figure 1.**
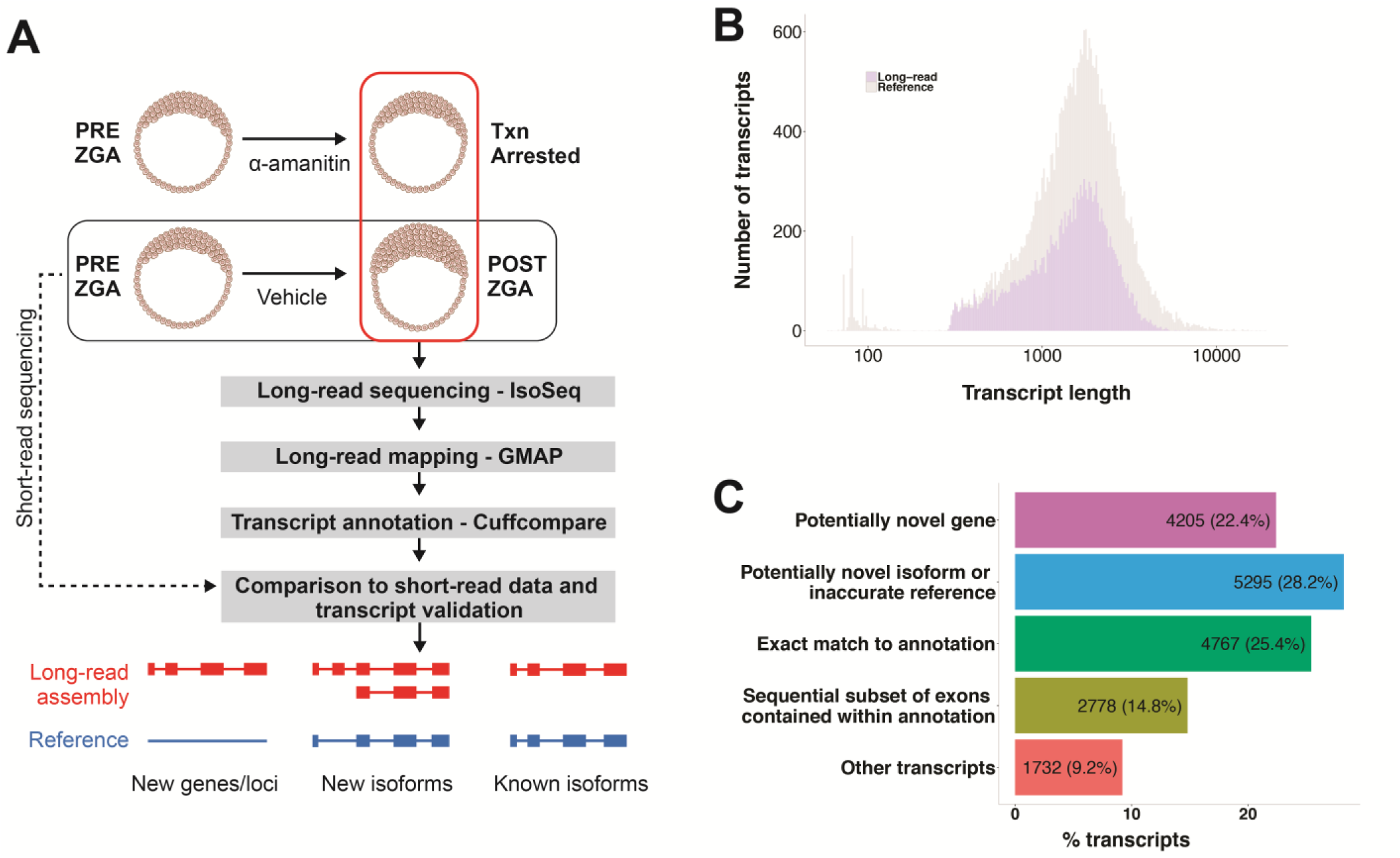
Overview of the embryonic zebrafish long-read transcriptome analysis. (A) Schematic of the long-read– based transcriptome reconstruction pipeline. Pooled RNA of α-amanitin / untreated embryos were collected and profiled with long-read sequencing. Temporally corresponding pre/post ZGA pooled embryonic RNA samples were profiled with short-read RNA-seq. Long-read raw data were assembled into transcripts using IsoSeq (O’Grady et al. 2016), mapped to the reference GRCz10 genome using GMAP (Wu and Watanabe 2005) and annotated against the reference transcriptome using Cuffcompare (Trapnell et al. 2012). Novel transcripts were compared to short-read data and computationally validated in constructing a final long-read augmented transcriptome. Select transcripts were experimentally validated. (B) Coverage of the reference GRCz10 zebrafish transcriptome by long-read sequencing. Shown in grey is a histogram of the number of RefSeq annotated transcripts along a range of transcript lengths. Shown in light purple is a similar histogram of long-read transcripts that overlap RefSeq annotations by any amount. 52.8% of reference transcripts have some overlap with the long read data. (C) Potential novelty in the long-read transcriptome. Long read data were compared for similarity in exon–intron structures against the reference annotation. A majority of the observed transcripts corresponded to potentially novel genes or isoforms.

We applied the IsoSeq pipeline (O’Grady et al. 2016) to cluster raw high-quality long reads into final assembled transcripts (see Methods), resulting in a combined 18,713 total non-redundant high-confidence isoforms from the treated and untreated samples. To provide a preliminary assessment of the zebrafish annotation representation in the long-read transcript dataset, we mapped the dataset to the most current version of the zebrafish reference genome (GRCz10) using GMAP, a mapper that supports gapped alignment (Wu and Watanabe 2005). 18,031 transcripts were successfully mapped to the reference. Only 3.6% of the dataset was unmapped, representing a very low percentage compared with unmapped zebrafish short-read data, which typically exceeds 20% (Aanes et al. 2014; Heyn et al. 2014). We then assessed overlap with the GRCz10 RefSeq annotation set consisting of high-confidence genes that exhibit conservation across multiple species (Pruitt et al. 2012). We selected RefSeq for this study because, while smaller in size than other catalogues such as Ensembl, it contains more accurate annotations and does not include incomplete transcripts that comprise a large proportion of Ensembl reference (Alamancos et al. 2015). Of the 15,159 RefSeq transcripts, 8,005 (52.8%) overlap with our long-read dataset (Fig. 1B). The coverage is fairly uniform for reference transcripts of lengths exceeding 200bp. Furthermore, the high percentage mapping to the reference sequence indicates the high quality of this dataset, and supports its use for identification of novel transcripts.

To assess potential novelty in the long read data, we first analyzed the long read transcripts for similarity in exon–intron structures with the RefSeq transcriptome (Fig. 1C). For each long-read transcript, we identified and categorized the most closely matching reference transcript using the Cuffcompare utility of the Tuxedo suite (Trapnell et al. 2012). 4,518 transcripts (24.1%) were found to be an exact match to the annotation. This low proportion of annotated transcripts identified in the data supports the presence of significant previously undiscovered transcriptional diversity. Interestingly, a recent human aggregate meta-transcriptome study reported a similarly low proportion of transcripts with annotations in the reference transcriptome (Iyer et al. 2015). A majority of the long read identified transcripts, 9,825 (56.5%) represent likely novel findings. 3,447 transcripts (21.6%) were potential novel transcribed regions (NTRs) and showed no overlap with the reference annotation. 5,558 transcripts (34.9%) were classified as potentially novel isoforms of previously annotated genes. This large group included transcripts with skipped exons, retained introns and other types of canonical alternative splicing events (Pan et al. 2008).

Consonant with our goal of accurate enhancements to the existing annotation, we were selective in which novel transcript types were studied. Two categories of transcripts for which eliminating sequencing and mapping artifacts was difficult, together comprising 23.7% of the data set, were therefore excluded from further analysis. Of these, 2,695 transcripts (14.4%) were classified as consecutive exonic subsets of the annotation, with missing exons that are present in reference transcripts. The missing exons were typically 5’ and/or 3’ end exons, whose absence in our long-read identified transcripts could have arisen through fall-off events of the active polymerase during sequencing or reverse transcriptase during cDNA preparation. A further 1,739 transcripts (9.3%) mapped to regions that are generally considered less likely to give rise to true transcripts. Included in this category were intronic regions, repeats and regions of exonic overlap on the opposite strand. While many of these transcripts could be artifacts of the sample preparation or sequencing process, they might also include potentially interesting findings (*see* Discussion).

### Characterization of Novel Transcribed Regions

Multiple analyses were performed to validate the novel transcribed regions detected by long-read sequencing. We first examined whether evidence of expression for these potentially novel transcripts were detectable in short-read data collected from RNA of embryos at comparable time points in development. After augmenting the reference transcriptome with the NTRs discovered by long-read sequencing, we mapped short-read data (see Methods) to the augmented transcriptome. An NTR example of a UCSC genome browser track is shown in Fig. S1A (Kent et al. 2002). This example illustrates that, with the augmented transcriptome, data from short read sequencing was successfully mapped to capture the full exonic structure of the novel transcript that is missing from the zebrafish annotation. To comprehensively characterize short read data support for the novel transcripts, we quantified the gene expression in these short read data against the NTR augmented transcriptome using *kallisto* (Bray et al. 2016). Of the more than two thousand largely non-overlapping novel transcripts discovered by long read sequencing in the untreated- and α-amanitin treated samples (see Figs. 2A, S1B), 89% and 86%, respectively, have support in the short-read data (TPM > 1). As evidenced in the quantified expression levels, many transcripts are highly expressed (median TPM = 3.5 and 4.2), and the overall exonic mapping rate of the short-read data improves with the augmented transcriptome in comparison with the RefSeq annotation from 68% to 85%.

**Figure 2.**
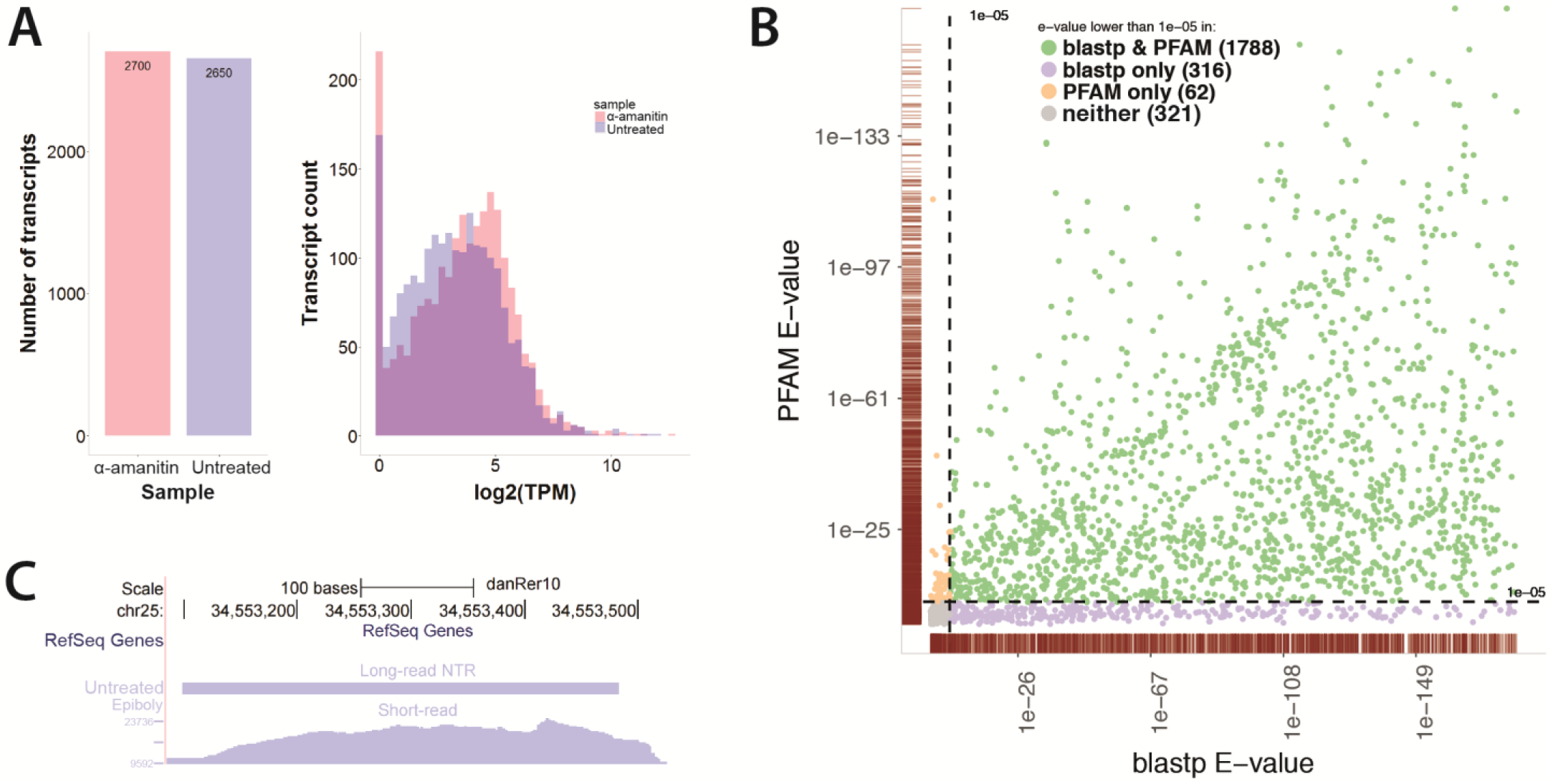
Characterization of novel transcribed regions (NTR) and coding potential. (A) Short read-support for all novel transcribed regions. Left panel: the numbers of long-read identified NTRs from the untreated and α-amanitin treated samples. Right panel: short read support for NTRs quantified by *kallisto* (Bray et al. 2016). Only 270 long-read transcripts in the α-amanitin treated condition and 343 in the untreated condition do not have short-read support (TPM < 1). Note that the these NTRs include both protein coding (see panels B, C and non-protein coding (see Fig. 3) transcripts. (B) Sequence homology and functional domain analysis. NTRs with high coding potential (determined by CPAT (Wang et al. 2013)) were tested for sequence homology with known proteins using *blastp* (Altschul et al. 1997), (x-axis) and for the presence of functional protein domains using hmmscan against the Pfam database (Finn et al. 2014), (y-axis). The color scheme indicates significance (e < 1e-05) by both *blastp* and PFAM (green), *blastp* only (purple), or PFAM only (orange). The gray markers in the lower left corner did not achieve significance in either analysis. The point densities in each dimension are indicated by the burgundy marginal rug plots along the x and y axes. (C) An NTR locus with homology to the human HIST2H2BE gene. Chromosomal location chr25:34,553,026-34,553,526 is shown using the UCSC genome browser. The empty top ‘ RefSeq Genes’ track indicates that no annotation exists in the reference. The middle track illustrates the novel transcript found by long-read sequencing in the untreated sample, and the bottom track shows corresponding mapped short reads after reference augmentation with the long-read detected NTRs. Note the high level of expression in the short-read data.

To identify novel transcribed regions that function as putative protein-coding genes, we analyzed these transcripts for evidence of their protein-coding potential, their conservation of sequence with known proteins and their function with known protein domains. We used CPAT (coding potential assessment tool) (Wang et al. 2013) to characterize the NTRs’ protein-coding potential. CPAT uses multiple metrics, including ORF length, ORF fraction of the sequence, Fickett’s score reflecting positional nucleotide usage in known coding sequences, and hexamer bias reflecting nucleotide dependencies in known coding sequences, to predict each sequence’s coding potential. Of the 4205 putative NTRs, CPAT identified 3255 sequences with high coding potential (*see* Methods). We evaluated 2487 of these high coding potential transcripts having a minimum ORF of 300 bp for sequence homology with known proteins using *blastp* (Altschul et al. 1997)] and for the presence of functional protein domains using *hmmscan* against the Pfam database (Finn et al. 2014). As shown in Fig. 2B, the vast majority were identified as encoding functional proteins using these assessments.

We considered the biological function of orthologs of a few selected novel protein-coding transcripts identified by our analysis. One interesting observation was the presence of novel orthologs of the histone variants of the human *H2AFX* and *HIST2H2BE* genes. Histones are particularly important in embryogenesis because of their requirement for cell proliferation during development. In both cases the zebrafish variants identified were very highly expressed in short read data as would be expected of histone genes (Fig. 2C and data not shown). *H2AFX* has an existing provisional gene zebrafish ortholog. Using long-read sequencing, we identified a novel *H2AFX* variant with a predicted ORF and a corresponding 127 amino acid peptide product that contains a 99.1% conserved histone core region. We found the previously annotated *H2AFX* ortholog expressed in the untreated sample, while the novel variant was expressed in the α-amanitin treated sample. In the case of the human *HIST2H2BE* gene, the orthologous novel gene we identified was not a previously known histone variant in zebrafish. This *hist2h2be* variant is present exclusively in the untreated sample and is therefore likely of zygotic origin.

We next considered the pool of NTRs that were not classified as protein coding genes by our analysis, and assessed whether they showed evidence of being evolutionarily conserved. It has been noted in previous studies that long non-coding RNAs (lncRNAs) exhibit moderate sequence conservation (Cabili et al. 2011; Derrien et al. 2012; Iyer et al. 2015). To assess the level of sequence conservation, we analyzed the transcripts using two measurements, the fraction of significantly conserved bases (phyloP algorithm (Pollard et al. 2010)), and the maximally conserved sliding window in a phylogenetic alignment of eight vertebrate species (phastCons algorithm (Siepel et al. 2005)). This analysis, similar to what was done to assess conservation in the human meta-transcriptome (Iyer et al. 2015), captures both independently conserved individual positions within a transcript and contiguous regions of high conservation (see Methods). We observed increased levels of conservation in 258 transcripts out of the pool of putative non-protein coding NTRs (24%) relative to random control regions (Fig. 3A). Specifically, 249 transcripts were classified as having higher base-wise conservation and 96 transcripts as having higher contiguous window-based conservation compared to random intergenic regions.

**Figure 3.**
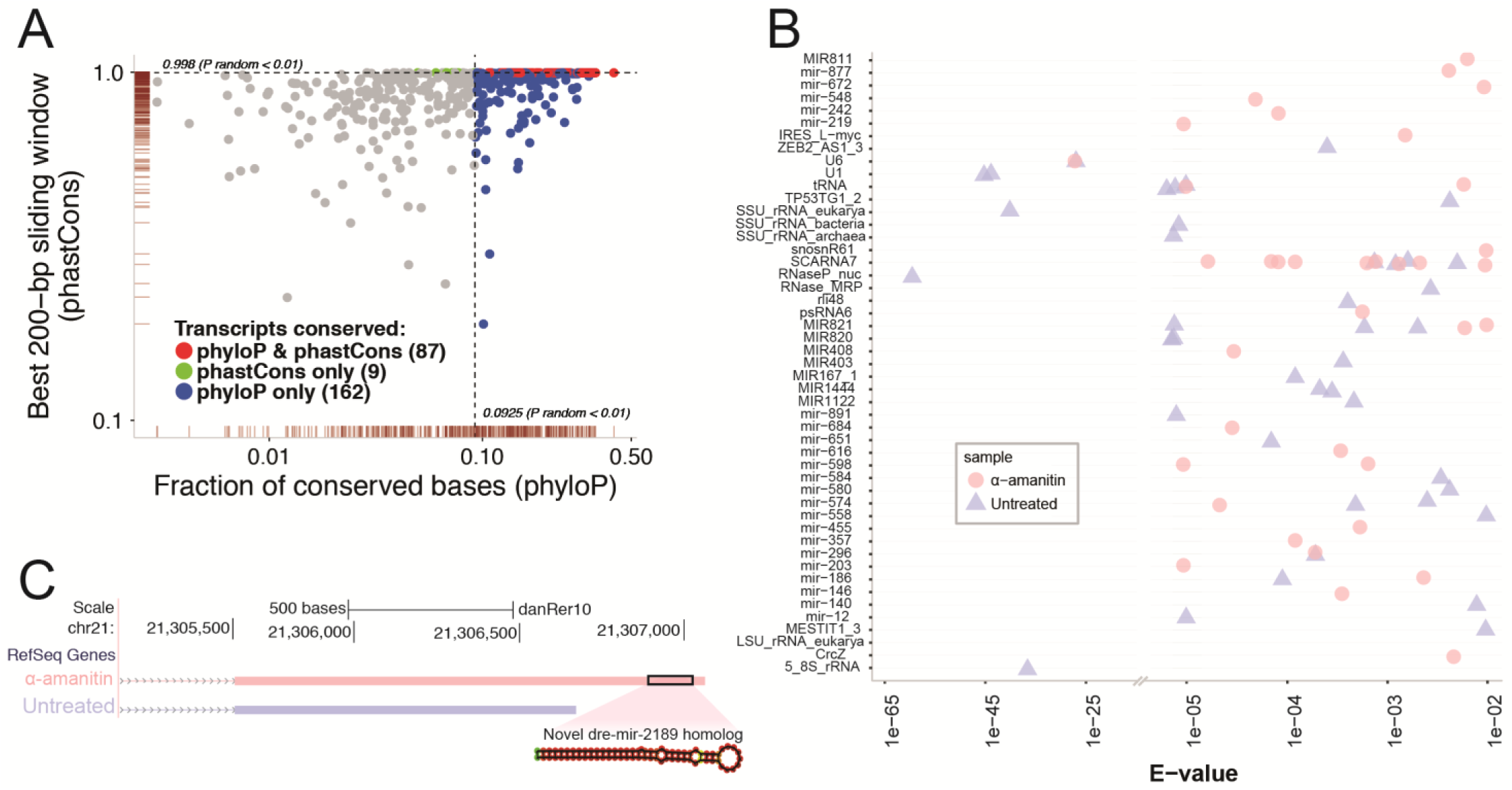
Characterization of putative non-coding NTRs. (A) Evolutionary conservation analysis of putative non-coding RNA NTRs. The scatter plot shows the base-wise transcript conservation levels (x axis) and the maximal 200bp window conservation levels (y axis). Base-wise transcript conservation levels were measured as the fraction of conserved bases (base-wise phyloP score > 1.5 (Pollard et al. 2010), see Methods for score selection). Window conservation levels were measured using a sliding window average PhastCons (Siepel et al. 2005) score across each 200bp region along the transcript. Blue points indicate transcripts that showed higher base-wise conservation (phyloP) relative to randomly selected intergenic regions (P_random_ < 0.01). Similarly, green points indicate transcripts with higher window-based conservation (PhastCons) relative to randomly selected intergenic regions (P_random_ < 0.01). Transcripts that met both conservation criteria are colored red. The marginal rugs along the x and y axes indicate the point density in each dimension. For clarity of presentation, data points with values of x < 1e-3 were omitted from the plot. (B) Functional analysis of putative non-coding RNA NTRs. The *cmscan* option of the Infernal tools (Nawrocki and Eddy 2013) was used to search for NTRs that were good matches to the Rfam database (Griffiths-Jones et al. 2003). Each point represents an individual transcript. Multiple transcripts can be mapped to the same non-coding RNA family. The points shown are above a default significance e-value (1e-02). The high abundance of relatively lower significance assignments (to the right of the break in the x-axis) largely represent miRNAs, which due to their short structures, have a limited maximal alignment significance. (C) Novel putative miRNA homolog. A novel transcript, containing a match to the miR-548 Rfam profile, was observed in the α-amanitin sample. An alternative isoform of this transcript with a shorter 3’ tail is observed in the untreated sample. The predicted miR match is located in the extended 3’ region, and appears to be a novel homolog of the existing zebrafish miR2189. The miR structure prediction was done using the Vienna RNA Websuite (Gruber et al. 2008).

Finally, we evaluated which non-protein coding NTRs had evidence of being functional non-coding RNA transcripts. We used the Rfam database (Griffiths-Jones et al. 2003), a collection of multiple sequence alignments and secondary structure profiles representing non-coding RNA families, and the *cmscan* option of the Infernal tools (Nawrocki and Eddy 2013) to search for NTRs that were good matches to the Rfam non-coding RNA families. Using the default significance threshold e-value of 1e-02, we identified 76 long-read transcripts that were matched to an Rfam profile (Fig. 3B). One particular NTR was found to be a match to the miR-548 Rfam profile. This transcript is specific to the α-amanitin sample, presumed to contain mostly maternally derived RNAs. An alternative isoform of this transcript with a shorter 3’ tail is observed in the untreated sample that consists of mostly zygotic RNAs. The predicted miR match is located in this extended 3’ region, and appears to be a novel homolog of the existing zebrafish miR2189 (Fig. 3C), as annotated by miRBase (Griffiths-Jones et al. 2008) that is not expressed in our samples. This pair of transcripts follow the prototype found in other studies (Mishima and Tomari 2016) showing that maternal transcripts possess longer 3’ tails as either putative targets or the miR structures themselves and has been implicated in the regulation of maternal to zygotic transition.

In order to generate a high confidence public resource of novel annotations, we applied stringent criteria. For protein coding NTRs, e-value of 1e-05 in either *blastp* against Uniprot or PFAM scan was required, leading to a total of 2487 novel protein coding annotations. For non-coding RNA NTR transcripts, correspondence of the results obtained by the conservation analysis and by assignment to non-coding RNA profiles identified 261 novel high confidence annotations. These newly annotated sequences will be of interest to investigate for their role in ZGA and early development.

### Characterization of Novel Isoforms for Annotated Gene Loci

We next studied the putative alternative isoforms observed in the long read data to determine which are true novel spliced isoforms of annotated zebrafish genes. For this analysis, we relied on the identification of valid exon-intron splice junctions to confirm alternatively spliced isoforms. The relatively sparse catalog of alternative splicing events in the RefSeq transcriptome makes it too limiting to use for validating the long read derived junctions. Instead, we relied on our high depth short-read RNA-seq data to detect splicing sites via reads that span across spliced exons. While identification of exact gene isoforms and their quantification from short-read RNA-seq data remains a very challenging problem, inference of splice junctions has recently become a tractable task for multiple spliced aligners (Engstrom et al. 2013). Therefore, we compared how well the splice junctions agree between our short and long-read data, and show the distribution of distances between the corresponding acceptor and donor sites (Fig. 4A). We inferred splice junctions in the short-read data using STAR (Dobin et al. 2013), and focused on the canonical (GT/AG) intron motif, which represents the absolute majority of both short and long read junctions. Of the approximately 33,000 such splice junctions with any intronic overlap between the short and long read data, 99% exhibit perfect correspondence for their donor and acceptor sites. This near perfect concordance of splice junction locations supports the accuracy of exon-intron boundary assignment from the long read data, and strongly supports the validity of the novel alternatively spliced isoforms reported.

**Figure 4.**
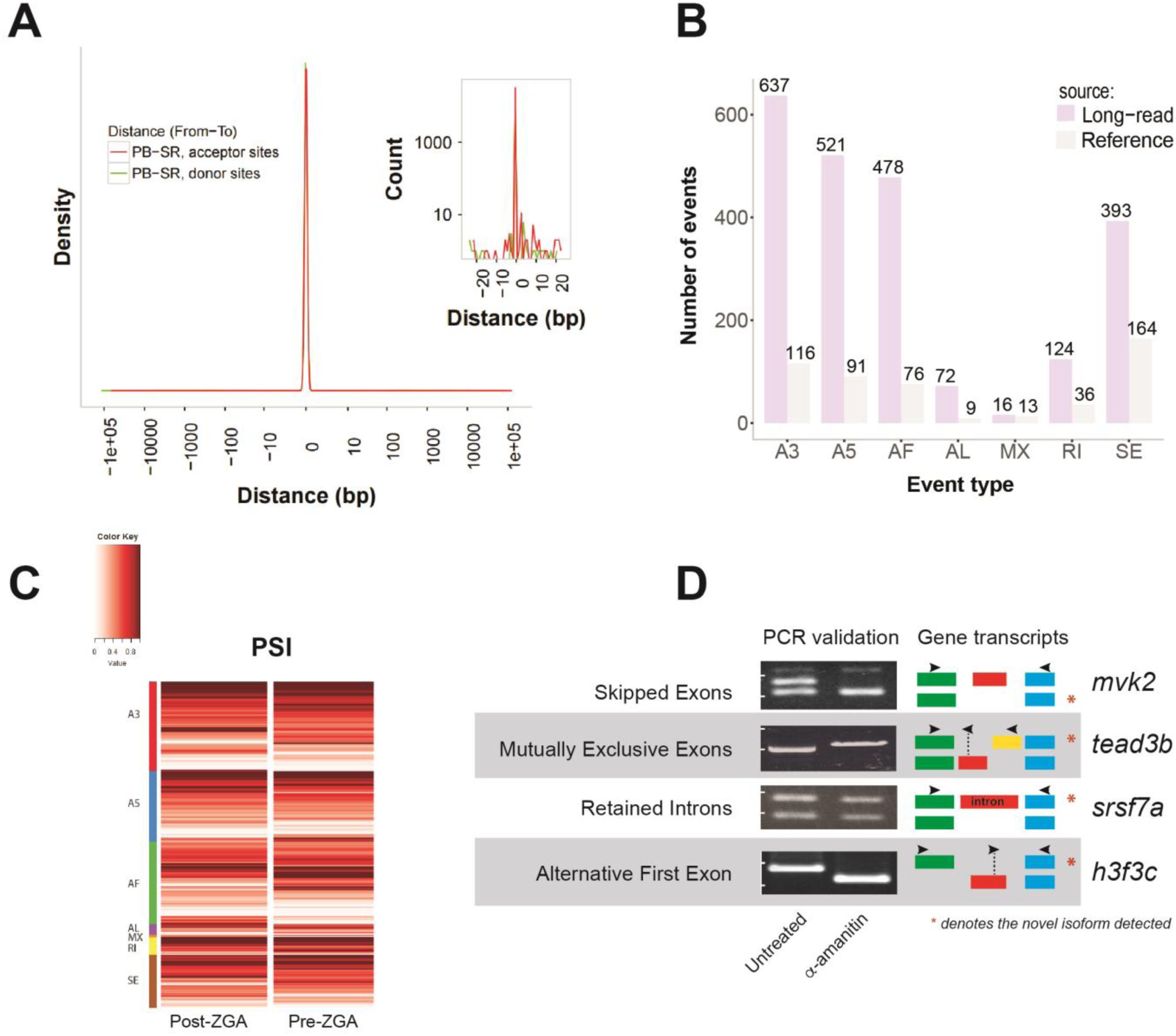
Characterization of novel isoforms for annotated gene loci. (A) Correspondence between long-read and short-read identified acceptor and donor sites in splice junctions. Shown in red is the distribution of distances from long-read determined acceptor sites to closest short-read determined acceptor splice sites. A similar histogram is plotted for donor sites in green, but is not well resolved due to superimposition of the plots. A zoom of the curves near zero is shown in the inset. Included in the plots are splice junctions for transcripts that have any overlapping introns as determined by long-read and short-read sequencing. Both curves peak heavily at zero indicating extraordinary agreement between the datasets. (B) Distribution of alternative splicing (AS) event types in the reference and long-read data. Alternative splicing events were identified using SUPPA (Alamancos et al. 2015). Listed AS include: skipped exon (SE), alternative 5′ or 3′ splice site (A5, A3), mutually exclusive exons (MX), retained intron (RI), and alternative first or last exon (AF, AL). (C) Percent spliced in index (PSI) (Schafer et al. 2015) for novel AS events from short-read RNA-seq data in early and late embryonic stages. PSI is plotted on a zero to one scale and indicates the fraction of the total transcript amount from each locus accounted for by a specific splicing event. The PSI values corresponding to each event in Panel B are plotted separately on each line and grouped by event type. For each novel AS event, PSI is shown from pre- and post-ZGA stage data. When assessed using Wilcoxon signed-rank test, FDR-corrected q < 0.05, the data show a significant increase in alternative 3’ untranslated UTR and in retained intron usage in the late embryonic stage. (D) Validation of novel splicing events in various AS categories using PCR reactions designed to distinguish isoforms. cDNA from α-amanitin / untreated samples was used for PCR. Primer sets, indicated by arrow (→,←) pairs, were designed to flank the splicing events, and the amplicon identity was confirmed by size. Exons are represented by colored boxes. For each novel AS event, expression of two isoforms was tested in each sample. The novel isoforms directly supported by PacBio reads are indicated by (*) for each pair of isoforms shown in the ‘ Gene model’ column.

In order to refine our list of putative novel isoforms, we quantified the number of alternative splicing events and compared the distribution of splice event types to that observed in the Refseq annotation (Fig. 4B). This quantification of exon skipping (SE), alternative 5′ and 3′ splice sites (A5/A3), mutually exclusive exons (MX), intron retention (RI), and alternative first and last exons (AF/AL) was performed using the SUPPA software package (Alamancos et al. 2015). While the reference annotation contains few alternative isoforms for known gene loci (1.08 on average), we discovered over two thousand novel alternative splicing events. The majority of these novel events were in the alternative 5′ and 3′ categories, reflecting alternative exon boundaries and the presence of multiple 5′ and 3′ UTRs. Although concerns have been raised about the resolution of PacBio sequencing in the 5’ end of transcripts (O’Grady et al. 2016), our reported novel splicing events are not affected by this potential technological shortcoming due to our requirement for the presence of valid donor and acceptor sites. Many alternative first exons were also identified, representing alternative transcription start sites (Fig. 4B). The alternative novel splice events enabled us to define a set of high confidence novel isoforms by including all long-read transcripts containing at least one such splice event. Our analysis identified a total of 1835 isoforms of previously known gene loci that were not annotated in the zebrafish RefSeq catalog, representing an approximately 50% increase in the number of isoforms identified per known gene locus. We used short read data to quantify the relative usage of the novel isoforms we have identified in samples from early and late embryonic stage samples. Measured as percent spliced in index (PSI), the ratio between reads including or excluding exons involved in each splicing event, the novel isoforms showed a significant increase in alternative 3’ untranslated UTR and in retained intron isoform usage in the late embryonic stage (Wilcoxon signed-rank test, FDR-corrected q < 0.05, Fig. 4C).

We next used exon-specific PCR reactions to validate examples for several splicing mechanisms underlying formation of novel isoforms detected in the long read data. The PCR primer selection that distinguishes the different isoform splicing mechanisms are shown schematically (Fig. 4D, see Supplementary Table 1). The following events detected in long read data were confirmed by PCR: skipped exon (*mvk2*), mutually exclusive exon (*tead3b*), retained intron (*srsf7a*), and alternative first exon (*h3f3c*). As detected in the long read data for these examples, the splice site usage seen by PCR differs in zygotically derived (untreated) and maternally derived (α-amanitin) samples. Therefore, our approach to identify novel isoforms of transcripts generated during the ZGA stage of development suggests that the current understanding of transcripts of important genes requires refinement.

### Novel Early ZGA Isoforms with Implicit Functional Significance

Included among the large number of novel isoforms expressed in early ZGA identified by our analysis was a long transcript encompassing repeats of the functionally important miR-430 microRNA. Heyn et al. established miR-430 as likely the first significantly-expressed zygotic transcript (Heyn et al. 2014). miR-430 links ZGA to the decline of the maternal program by specifically targeting a significant proportion of all maternal transcripts for silencing and degradation (Giraldez et al. 2006). Although miR-430 is known to be present in early ZGA, it had not previously been recognized as an excision product of a contiguous transcript.

We inspected the miR-430 cluster in order to observe whether additional transcripts arise from this region during early ZGA. Unexpectedly, we observed a novel 4-exon transcript spanning 9kb of genomic sequence overlapping 22 miR-430 repeats, which we refer to as “mega-miR-430”. (Fig. 5A). Notably, a large fraction of miR-430 repeats occurred within mega-miR-430 introns. As mega-miR-430 was not detected upon α-amanitin treatment, it appeared to be purely of zygotic origin. Confirmatory qPCR assays for two unique mega-miR-430 regions showed its level was dramatically reduced by α-amanitin treatment (Fig. 5B). We assayed miR-19a levels as a control, given its exclusive expression during ZGA (Wei et al. 2012), and found a comparable reduction in its expression with α-amanitin treatment (Fig. 5B). The capacity to generate multiple copies of miR-430 from each mega-miR-430 transcript (see Fig. 5C) probably contributes to the high miR-430 levels reported during early ZGA (Takacs and Giraldez 2016). The generation of mature, functional miRNAs from the introns of larger transcripts has been previously recognized (Westholm and Lai 2011). We propose that mega-miR-430 gives rise to several functional miRNA units per transcript by intron splicing and subsequent miRNA processing. If mega-miR-430 is transcribed by RNA Polymerase II in parallel to polymerases generating pre-miRNAs from individual miR-430 genes, this could contribute to the high levels of miR-430 transcript observed at the start of ZGA. In comparison with most other developmentally-regulated genes (see Supplemental Fig. S2), we find that miR-430 is extraordinarily sensitive to RNA polymerase II inhibition by α-amanitin, which may reflect the fundamental role miR-430 plays during ZGA in zebrafish. These data suggest that important miRNAs have a complex processing strategy that contributes to a range of isoforms that may have different activities and targets.

**Figure 5.**
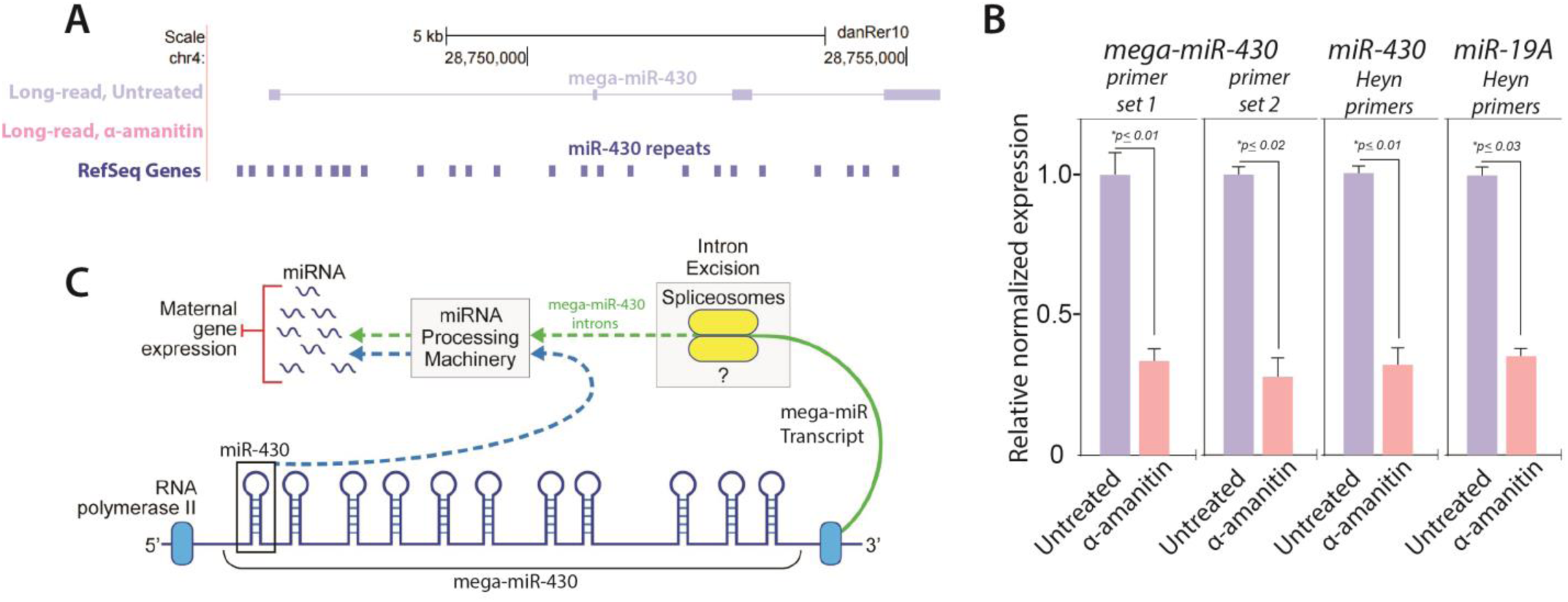
Novel validated transcript encompassing miR-430 repeats. (A) A novel transcribed region locus encompassing the miR-430 repeats. The top panel shows genomic location chr4:28,745,803-28,757,055 with annotated miR-430 repeats in the reference. The two lower panels illustrate that the novel transcript found by long-read sequencing, which encompasses the miR-430 repeats, is specific to the untreated sample. (B) Experimental validation of mega-miR-430 transcript in zebrafish embryo RNA by RT-qPCR. The same RNA samples used for long-read sequencing were assayed by RT-qPCR to experimentally validate mega-miR-430 and relate this validation to the validated behavior of established transcripts. mega-miR-430 transcript was assayed using two independent primer sets spanning different introns. Both sets give nearly identical results and show susceptibility of the transcript to α-amanitin, agreeing with long-read sequencing data. Prominent early-onset “zygotic-only” genes miR-430 and miR-19a also show high abundance and strong susceptibility to α-amanitin using previously reported primers and consonant with a previous report (Heyn et al. 2014). All transcripts measured by qPCR were normalized to the *Hprt* mRNA, observed to be unchanged during treatment. (C) Proposed functional mechanisms of mega-miR-430 transcription. Mega-miR-430 serves to enrich miR-430 transcription by encoding a long single transcript that contains miR repeats in its introns. The generation of mature, functional miRNAs from the introns of larger transcripts is an established mechanism (Ramalingam et al. 2014). By this mechanism mega-miR-430 gives rise to several functional miRNA units per transcript after intronic splicing and miRNA processing. If mega-miR-430 is transcribed by RNA Polymerase II in parallel to polymerases generating pre-miRNAs from individual miR-430 genes, this would amplify the copies of individual miRs through a single larger transcript, enabling the abundant levels of miR-430 transcript observed at the start of ZGA. Illustration by Jill Gregory (Mount Sinai Health System).

## DISCUSSION

In this study, we developed an analysis pipeline that leverages current long-read sequencing technology to greatly improve the resolution of the zebrafish transcriptome annotation. We focused on early zebrafish development due to its diverse transcriptional repertoire, and because zebrafish have unique attributes for a vertebrate model that allow detailed study of how the zygotic genome becomes activated. We combined the ease of using the well-established transcriptional inhibitor α-amanitin (Lee et al. 2004) to identify transcripts that are derived from the zygotic genome. We identified thousands of previously unknown presumptive protein coding transcripts, as well as thousands of novel isoforms of previously annotated genes. Our systematically validated findings represent a greatly expanded transcriptome annotation for the research community.

We identified and validated more than 2700 new, previously unannotated transcripts and nearly two thousand new isoforms of previously known gene loci that are not annotated in the zebrafish RefSeq catalog. The newly annotated loci represent a 20% increase over the latest RefSeq catalogue, and the novel isoforms are an approximately 50% increase in the number of isoforms identified per known gene locus. The novel transcribed regions were stringently validated using short read sequencing data, sequence homology and functional domain conservation. For validation of the novel isoforms, we found a near perfect concordance of splice junction locations using short read data that supports the accuracy of novel exon-intron boundary assignments from the long read data. Our analysis of short read data finds that the isoforms identified using long read data show a significant increase in two types of alternative splicing events. Both alternative 3’ UTR isoform usage and retained intron usage are significantly increased in samples from late stage versus early stage embryos (see Fig. 4C). Our analysis of novel 3’UTR isoforms usage is consonant with the study by Heyn *et al*. of previously identified zebrafish splice events (Heyn et al. 2014). The increase in retained intron usage seen in late stage embryos is notable given the importance of this mechanism to introduce regulatory sequences, such premature stop codons, during cell differentiation (Pimentel et al. 2016).

Examination of various novel transcripts support the utility of our new annotation. For example, we identified two novel orthologs of the histone variants of the human *H2AFX* and *HIST2H2BE* genes, which are noteworthy given the importance of histone subtypes in cell division during ZGA. We also identified many novel miRNA precursor isoforms, including a novel homolog of miR2189 and a previously unrecognized transcript containing 22 miR-430 repeats. miR-430 has a critical role in silencing maternal transcripts (Giraldez et al. 2006). Our study has identified, for the first time, the mechanism of production of the miR-430 family from intronic regions of a mega-miR-430 contiguous transcript. We also confirmed the existence of specific novel alternative splicing events inferred from the long read data via PCR experiments, including skipped exon, mutually exclusive exon, retain intron and alternative first exon (see Fig. 4D).

The long read technology used has the limitation of being insensitive to mature miRNAs, as very short sequences are excluded (Rhoads and Au 2015). In addition, while we identified nearly 2500 validated novel protein coding RNAs, we found only a few hundred novel ncRNAs. The RNAs in the libraries utilized were poly(A) tail enriched. As ncRNAs are less likely to be poly(A) tailed (Pauli et al. 2012, Chew et al. 2013), this selection bias may have contributed to their under representation. The relatively low number of novel ncRNAs in our final annotation resource may also be the result, in part, of our validation pipeline. Coding RNAs are more likely to show secondary structure and sequence pattern recognized by computational validation methodologies. In contrast, ncRNAs lack these computationally recognizable features. Furthermore, many transcripts were excluded from analysis due to their containment in intronic regions, repeats and regions of exonic overlap on the opposite strand. Such transcripts excluded from analysis may be relatively enriched in ncRNAs (Pauli et al. 2012). In the development of our analysis pipeline, we accepted less comprehensive reporting of novel transcriptional annotations in pursuit of the overall goal of generating a highly reliable expanded annotation resource.

## METHODS

### Zebrafish treatment

Zebrafish were maintained according to standard protocols and embryos were obtained during natural spawning of either ABNYU or TAB14 WT adults. The IACUC committee of the Icahn School of Medicine at Mount Sinai approved all protocols. Adult wild-type ABNYU and TAB14 zebrafish were maintained on a 14:10 light:dark cycle at 28°C with feedings twice daily. Fish were removed from the maintenance system and placed into mating tanks within 1 hour of light cycle activation. Fish were allowed to mate spontaneously and returned to sex-segregated maintenance tanks by net capture. Fertilized eggs were collected by net and incubated at 28°C in egg water (0.6 g/L Crystal Sea Marinemix from Marine Enterprises International, Baltimore, MD, dissolved in dH_2_O) containing methylene blue (0.002 g/L) All quantitative PCR (qPCR) studies were conducted from total RNA isolated from three independent biological replicates. The procedure for achieving suppression of ZGA is treatment of one-to-four-cell embryos by injection of 0.2nmol of the RNA Polymerase inhibitor α-amanitin. Pools of 15 embryos each for both untreated wild-type embryos and embryos injected with 0.2nmol α-amanitin to abolish zygotic transcription, as described previously (Zamir et al. 1997). Fertilized eggs were collected and staged based on morphological criteria to identify embryos at pre (256-cell stage) and post-ZGA (6hpf).

### RNA Isolation, Reverse Transcription, and RT-PCR

RNA was isolated from zebrafish embryos via standard Trizol protocol as described (Kent et al. 2016). Reverse transcription was accomplished using the Superscript III kit from Invitrogen with 2ug total RNA. RT-PCR reactions were prepared with 50ng cDNA per reaction and RedTaq reverse transcriptase mix (Sigma). Annealing temperature was 58° C and extension time was 3 minutes. RT-qPCR reactions were prepared with 20ng cDNA per sample and set up according to the Promega GoTaq 2-Step SybrGreen™ kit using a fast 2-step protocol on the Agilent Mx3000P qPCR system and initial melting time of 3 minutes. Dissociation curves were also obtained. All measurements from a minimum of 4 replicates were normalized by delta-delta C*t* to the housekeeping *Hprt* gene. Measurements for each set of technical replicates (untreated, α-amanitin) were then normalized to the median of the untreated sample. Error bars represent standard error. Gel analysis of PCR in Fig. 4D were performed on 1% agarose gel or 10% polyacrylamide gel after 25-40 cycles. Primer sequences used are included as part of the Supplemental Materials.

### Short-read RNA sequencing data and analysis

For short read sequencing, RNA was isolated from sample sets of approximately 20 embryos at 2.5 hours post fertilization (hpf) and 5.25hpf stages. RNA quality was assessed by Agilent Bioanalyzer. Samples with a RIN score of 9 or greater were used to generate cDNA libraries following treatment with the Ribo-Zero rRNA removal kit (Illumina).Tru-Seq cDNA libraries were generated and sequenced on the Illumina Hi-Seq 2500 platform (100 base pair paired-end). Resulting library sizes exceeded 25M paired reads. QC was done using the *FastQC* package. RNA-seq alignment to the GRCz10 RefSeq *Danio rerio* reference and splice junction identification was done using STAR (Dobin et al. 2013). Isoform quantification was done with *kallisto* (Bray et al. 2016).

### Single Molecule Real-Time Sequencing (SMRT-Seq)

*cDNA preparation*: Total RNA from 256-cell stage embryos was collected for PacBio SMRT library generation. RNA quality was assessed by Agilent Bioanalyzer, and samples with RIN scores of >9 were retained. RNAs were poly-A selected using oligo-dT beads. First-strand cDNAs were generated and amplified using the Clontech SMARTer PCR cDNA Synthesis Kit. The resulting cDNA libraries were purified using 0.6× volume Agencourt AMPure PB Beads (Beckman Coulter, Life Sciences Division), as specified by the supplier (Pacific Biosciences). Libraries were then size-selected using the BluePippin system (Sage Science, Inc.) into four separate bins each: <1kb, 1–2 kb, 2–3 kb, and >3kb. *Library construction and sequencing:* SMRTbell libraries were prepared from 0.5 and 1.0 µg of each cDNA size class using the PacBio Large Insert Template Library Prep Kit. The samples were sequenced on the PacBio RS using “C2” chemistry. SMRTcells were loaded using a combination of diffusion and MagBeads. Two to four SMRTcells were used for each size bin, depending on the estimated relative abundance of transcripts in that size range. *Primary Data Analysis*: Long reads produced by the PacBio RS sequencer were processed via the PacBio IsoSeq pipeline to generate full-length refined consensus transcripts. In particular, sequences containing both 5′ and 3′ adapters were identified, and the adapters and poly-A/T sequences were trimmed. The retained high-QV (high quality) isoforms were aligned to the zebrafish genome assembly GRCz10 using the splice-aware mapper GMAP (Wu and Watanabe 2005). Identical transcripts with minor 5’ differences were merged using the ‘ 5’ collapse’ option.

### Characterization of long-read data with respect to the existing Zebrafish annotation

We used the Cuffcompare utility of the Tuxedo suite (Trapnell et al. 2012) to categorize each long-read transcript with respect to its most closely matching reference transcript. The Cuffcompare class codes underlying long read transcript classification in Figure 1D were: ‘u’ for ‘Potentially novel gene’; the set of ‘ e’ and ‘j’ for ‘Potentially novel isoform or inaccurate reference’; ‘=‘ for ‘Exact match to annotation’; ‘c’ for ‘Sequential subset of exons contained within annotation’; and the set of ‘i’,’o’,’p’,’r’,’s’,’x’ for ‘ Other transcripts’. The code ‘ e’ (88 transcripts), defined as ‘ Single exon transfrag overlapping a reference exon and at least 10 bp of a reference intron’ was added to the ‘Potentially novel isoform or inaccurate reference’ category due to potentially inaccurately defined exon boundaries in the existing Zebrafish transcriptome.

### Assessing short-read data support for Novel Transcribed Regions (NTR)

To assess whether independently generated short-read data contains evidence for the newly discovered putative transcripts, we first augmented the reference transcriptome with gene models for the corresponding set of NTRs using Cuffmerge (Trapnell et al. 2012), and then estimated the short-read transcript abundance against the augmented transcriptome with *kallisto* (Bray et al. 2016).

### Characterization of NTRs with high protein-coding potential

We first classified the entire pool of NTRs with respect to their protein-coding potential (CP) using CPAT, v1.2.2 with the threshold parameter previously estimated for zebrafish (Wang et al. 2013). The NTRs with high CP were analyzed for conservation of sequence with known proteins and conservation of function with known protein domains. TRANSDECODER, v.3.0.0 (Haas et al. 2013), was used to extract ORFs from the high CP transcripts. The obtained ORFs were matched against Uniprot (release 2016_06) using *blastp*, v2.2.26, and scored against PFAM-A (release 30.0) using *hmmer/3.1b1*.

### Characterization of putative non-coding NTRs

*Evolutionary conservation analysis:* The evolutionary conservation of the putative non-coding NTRs was studied using two measurements, the fraction of significantly conserved bases (phyloP algorithm (Pollard et al. 2010), and the maximally conserved 200 nt sliding window (PhastCons algorithm (Siepel et al. 2005)) in a phylogenetic alignment of eight vertebrate species. To control the false discovery rate, we also measured the conservation of non-transcribed regions using these metrics by random sampling of contiguous length-matched intervals. The sampled non-transcribed regions were defined to exclude sequences containing known genes, extended by +/- 1Kb flanking regions. For both base-wise conservation (phyloP) and contiguous window conservation (PhastCons) analyses, cutoffs for significant transcripts were determined by setting the false discovery rate (FDR) to 0.01, i.e. controlling the rate of observing elements with equivalent conservation scores within non-transcribed sequence space at a level of 0.01. At this FDR level, the phyloP average transcript conservation score was 0.0925 (9.25% of transcript bases conserved at phyloP score > 1.5). The phyloP score was chosen as the 95^th^ percentile of all scores in the non-transcribed randomly selected sequence space. For sliding window conservation, FDR < 0.01 corresponded to an average PhastCons probability of 0.998. *Matching known RNA families in RFAM:* A search for matches to known noncoding RNAs families, within the pool of NTRs with low CP, was performed using the *cmscan* option of Infernal v1.2 (Nawrocki and Eddy 2013) against the pre-calibrated covariance models of the Rfam database release 12.0 (Griffiths-Jones et al. 2003), keeping default *cmscan* settings and discarding predictions with e-value > 1e-02.

### Long-read splice site accuracy estimation

To assess the accuracy of splice sites in ISO-seq transcripts we computed the distance distribution for donor/acceptor sites between all the Iso-seq transcripts and the short-read used in the study. First, each Iso-seq exon junction was examined for canonical splice acceptor (AG|N) and splice donor sites (N|GT). Second, each Iso-seq junction with its corresponding genomic positions was compared against the pool of junctions derived from Illumina short-read sequencing (*as described* in Methods section “*Analysis of short-read RNA sequencing data*”). Short-read splice junctions identified by STAR were split into groups in accordance with the intron-motif nomenclature used by STAR and only the subgroup representing the canonical junctions was retained. Finally, the distances from acceptor and donor splice sites in the Iso-seq data to nearest corresponding sites in the short-read data were computed.

## DATA ACCESS

The long-read and short-read RNA-seq data generated for this study have been deposited to GEO (accession number: GSE101843), and private reviewer links have been created. The curated resource resulting from this study, including all computationally validated novel transcribed regions and gene isoforms, is available at <u>http://slapp04.mssm.edu/download/zebrafish-enhanced.gtf</u>.

## ACKNOWLEDGEMENTS

We acknowledge Dr. Side Li for performing the PCR experiments and Christopher Smith for help with figure design. All sequencing services performed were supported through NIH grants 5R01CA154809 and 5R01HL103967. The development of the analysis pipeline was supported by NIH grant U19 AI117873.

